# Rapid Electro-optic Two-photon Bessel Light Sheet Fluorescence Lifetime Imaging for Tissue Volume Interrogation

**DOI:** 10.64898/2026.11.03.763887

**Authors:** Prashant Kumar, V. Rose Knight, Daniel Elliott Lysko, Jung-Gun Kim, William S. Talbot, Adam Bowman, Mary Beth Mudgett, Soichi Wakatsuki, Mark Kasevich

## Abstract

We report a two-photon light sheet electro-optic fluorescence lifetime imaging microscopy (TPLS-EO-FLIM) technique based on an electro-optic (EO) modulation detection module. Scanning two-photon Bessel light sheet excitation was used to achieve in-depth volumetric imaging. The low cross-section of multiphoton excitation requires high detection efficiency, and electro-optic detection enables fast wide-field FLIM with a scientific complementary metal–oxide–semiconductor (sCMOS) detector. Phototoxicity is minimized by utilizing signal from the entire illuminated sample area in a wide field camera exposure, and time-resolved volume acquisitions are feasible in minutes, with cross-sectional field of view and light sheet length over 300 microns. The method has sub-nanosecond lifetime resolution and is capable of accurate FLIM images in 10 ms exposures with as low as a few hundred photons per pixel. The technique was demonstrated through volumetric FLIM of a green fluorescent protein (GFP) labeled zebrafish embryo heart, of multi-fluorescent protein (multi-FP) labeled plant root structures and of dynamic cardiac cycles in live zebrafish embryos. For the latter, we recorded the cardiac cycle dynamics of a zebrafish embryo at 50 frames per second and measured biomechanical parameters such as heart rate, ventricle/atrium areas, atrioventricular valve dynamics, and fractional area change. Taken together, these studies show the system’s capability for rapid, high-resolution, and functional imaging in diverse biological contexts.

## Introduction

Fluorescence lifetime imaging microscopy (FLIM) probes the fluorophore environment. It provides insights into parameters such as pH, molecular concentration, and the presence of other molecules. It has been widely used in biological sciences for various applications such as drug discovery, biophysical analysis, and biochemistry [1, 2]. Despite its utility, FLIM imposes challenging constraints on detector performance, especially in the context of quantum efficiency, temporal resolution, and field of view (FOV). Time-correlated single photon counting (TC-SPC)-based lifetime measurement is a state-of-the-art method [1, 3]. However, the low quantum efficiency of highly time-resolved detectors limits it for low photon count applications, and single-pixel detection requires scanning, which makes it slow for extensive FOV interrogation. The advances in fast multi-pixel detectors, such as single photon avalanche diode (SPAD) arrays [4–6], silicon photomultipliers (SiPM)[7–15] and superconducting nanowire single-photon detectors [16–19], can be helpful for wide-field FLIM, but their development is still in progress [20–24].

A recent advance in fast detection methods is electro-optic (EO-) modulation-based wide-field detection, developed by Bowman et al. [25, 26]. EO-FLIM enables simultaneous whole-FOV acquisition on standard scientific cameras by encoding time resolution in the polarizations of fluorescence emission photons, which can then be split between optical gates. It combines the required temporal resolution to facilitate FLIM measurements with the high quantum efficiency of state-of-the-art scientific cameras to enable efficient lifetime detection. The resonant EO-FLIM method has been demonstrated for single-molecule [26] and kilohertz *in vivo* [27] imaging, as well as compact configurations sensitive to multiple lifetimes per pixel [28]. Pockels-cell-based FLIM has also been realized in time-gated rather than resonant configurations [29].

On the other hand, light sheet fluorescence microscopy (LSFM) has emerged as a promising technique for fast volumetric interrogation of biological specimens. Resonant light sheet EO-FLIM has been recently demonstrated, enabling lifetime-resolved imaging of 0.32 mm^3^ volumes in less than two minutes [30]. The light sheet method accelerates biological research, spanning from study of organelles to multicellular organisms, through its various specimen compatible excitation-detection geometries [31–39]. Light sheet configurations with cameras parallel to the sheet acquiring whole frames simultaneously minimize phototoxicity because signal is gained from all parts of the sample on which light is incident.

Two-photon excitation is attractive for many reasons. Excitation probability becomes quadratic in intensity rather than linear as in the single-photon case, meaning that excitation and damage is far more likely to occur only in the plane of the laser focus. Infrared light also scatters much less than visible light in biological tissues, allowing deeper tissue penetration. Multiphoton excitation confers benefits of UV excitation, including photoactivation, photolysis, and access to a broader range of fluorophores, without the global photodamage, absorption by endogeneous chromophores, and constraints on excitation optics that make UV excitation challenging. Historically, two-photon excitation has been limited by a photon budget that demands photon-efficient detection, as well as its voxel-at-a-time acquisition speed [40, 41]. Previous studies have combined two-photon and light sheet microscopies to demonstrate penetration depths of several hundred microns [42], or twice as much as single-photon excitation [43], substantial speedups compared to scanned two-photon microscopy [43, 44], and increased field of view of Bessel light sheets [45]. Two-photon light sheet fluorescence microscopy has enabled imaging of samples with high-scattering tissue such as hatched fish [46], 4D reconstruction of a live zebrafish heart [47] and whole-brain imaging [48], where infrared illumination prevents the accidental excitation of blue photoreceptors. Two-photon light sheets enable a wider and more homogeneous region of high-quality field of view than their single-photon counterparts due to reduced scattering and the suppression of nonlinear excitation in the diverging edges of the sheet, and light sheet methods can offer more time-efficient acquisition of three-dimensional multiphoton datasets [43].

Here, we report the development of a two-photon Bessel light sheet fluorescence lifetime microscope (TPLS-EO-FLIM) based around electro-optically modulated detection via a Pockels cell [27]. The combination of two-photon Bessel LSFM with electro-optic detection enables rapid, volumetric acquisition of FLIM data with minimal phototoxicity, making it well-suited for live and deep tissue imaging. As we show below, this integration allows dynamic mapping of biological systems across large 3D volumes.

We first validated the technique using non-biological reference standards—fluorescent beads and a fluorescent calibration slide. To demonstrate biological applications, we imaged a live zebrafish embryo heart and recorded its heartbeat at 50 frames per second with spatial and temporal lifetime information, and imaged multiple fluorescent probes with contrasting lifetimes labeling proteins in *Arabidopsis thaliana* seedlings. The cardiac dynamical imaging led us to estimate the various biophysical parameters such as heart rate, ventricle and atrium areas, atrioventricular valve dynamics, and fractional area change.

## Results

The implementation of TPLS-EO-FLIM is shown in Figure 1 and Fig. S1. The geometry of the light sheet was characterized by exciting a fluorescein solution. Its length was observed to be *∼* 340 *µ*m with a 20X illumination objective (Supplementary Information 2). This length covers most of the 4X field of view. The scan volume of the sheet depends on the scan range of the galvo-scanner and the back aperture of the objective lens and was kept to be *∼* 400 *µ*m during the experiments. The thickness of the light sheet was measured to be *∼* 2.5 *µ*m. The field of view was circular, with diameters of *∼*176 *µ*m and *∼*376 *µ*m for the 10X and 4X detection objectives, respectively. The corresponding resolution of the imaging system was observed to be 1.96 *µ*m and 3.0 *µ*m, respectively (Supplementary Information 5). These values are larger than the diffraction-limited values (*∼*0.5 *µ*m and *∼*1.3 *µ*m), owing to the finite bead size and residual aberrations in the detection path.

**Fig. 1.**
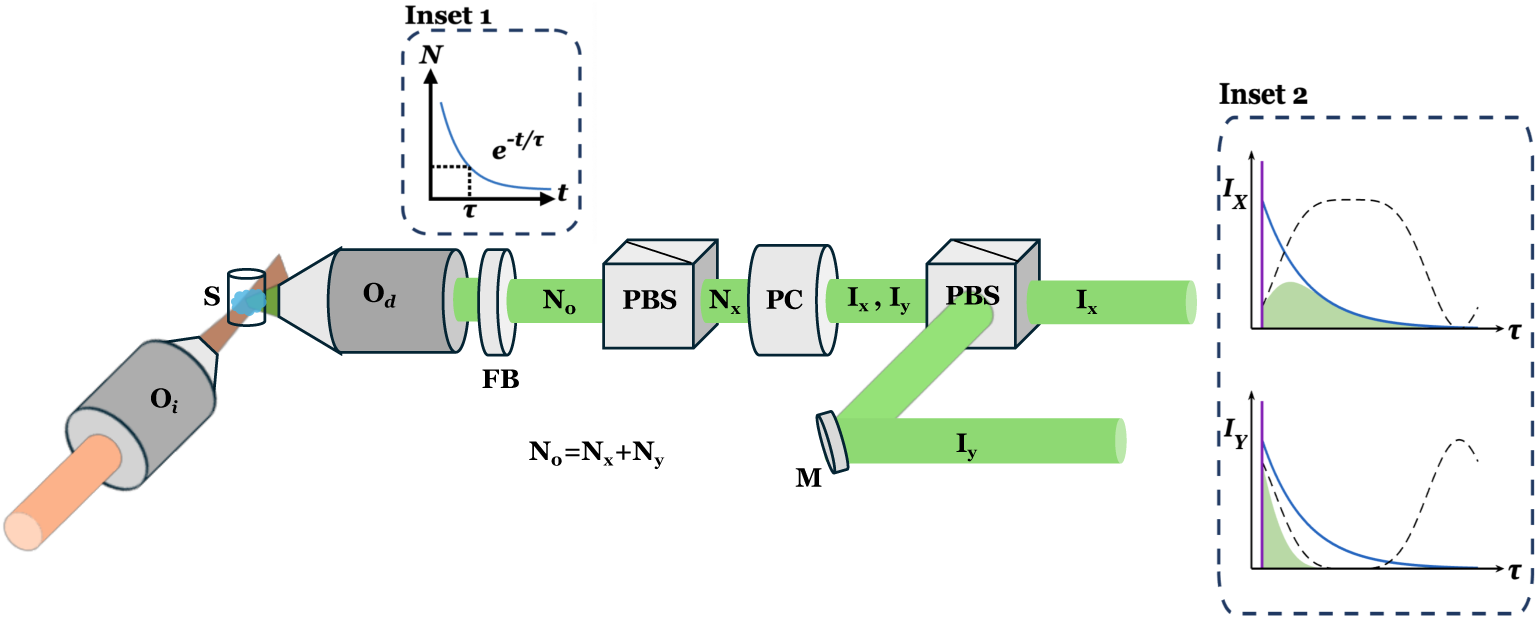
Schematic diagram of Electro-optic imaging based wide-field two-photon light sheet FLIM: The illumination and detection subsystems are arranged orthogonally for two-photon digitally scanning light sheet interrogation. A 100 fs pulse laser operating at a 930 nm wavelength with a 39 MHz repetition rate produces a digitally scanned two-photon Bessel light sheet through an illumination objective (*O_i_*, 20x/0.40) for specimen (S) sectioning (Fig. S1). The specimen is translated in the XYZ directions for interrogation by the light sheet. Fluorescence emitted from the specimen is collected by a detection objective (*O_d_*), filtered through a filter box (FB) containing long-pass and notch filters, and then directed to a PBS-Pockels cell-PBS combination for time stamps as a function of applied voltage (V) on the Pockels cell (PC). The fluorescence light is split into its time-binned X and Y polarized components (*Ix*, *Iy*), which are then focused onto an sCMOS camera (shown in the Fig. S1) through a tube lens. The PC is synchronized with the laser repetition rate at 39 MHz. Inset 2 presents a cartoon of the laser excitation (purple) and Pockels cell gating (dotted) with the exponential decay (blue), and the resulting detected intensity (green) at the camera, where a single camera exposure integrates over millions of the synchronized laser and RF cycles. Multiple cycles are depicted in Fig. S1, and the phase difference between the laser excitation and gating function depicted can be varied. See Methods section and Supplementary Information 1 for a detailed discussion of the setup. Abbreviations: *I_x_, I_y_*, intensity associated with x-polarized and y-polarized states; *N*_0_*, N_x_, N_y_*, photons associated with unpolarized, x-polarized, and y-polarized fluorescence; PBS, polarizing beam splitter; *τ*, fluorescence lifetime; M, mirror.

For lifetime imaging, the Pockels cell was driven with a 19.5 MHz sinusoidal voltage. A laser pulse was picked at a 39 MHz repetition rate to synchronize with the Pockels cell modulation. The lifetime instrument response function (IRF) was recorded by sweeping the phase delay between the Pockels cell drive and the laser pulse excitation of a quenched fluorescein solution. The quenched fluorescein solution has an extremely short lifetime (on the scale of picoseconds; see Supplementary Video 1 and Supplementary Information 3).

### Fluorescence lifetime demonstration of fluorescent beads

Yellow-green (1.0 *µ*m) and Nile Red fluorescent beads (1.0 *µ*m) were imaged as a reference sample to assess the system’s lifetime-based contrast performance [49]. The mixture of beads was embedded in agarose gel and confined within a rectangular glass capillary attached to the scanning stage. A two-photon scanning light sheet was employed for excitation. The exposure time was set to 50 ms per frame. The sample stage was systematically moved in 1.0 *µ*m increments to achieve volumetric scanning. Fluorescence emission was collected using a 10X/0.50 NA objective. Fluorescence lifetime was determined from the ratio of gated and ungated images at a fixed phase delay between the Pockels cell drive and the laser pulse (see Methods section).

Analysis showed distinct mean lifetime values of approximately 4.9 ns for yellow-green beads and 3.1 ns for Nile Red beads (Figure 2), validating the system’s potential to resolve fluorophores based on their unique fluorescence decay profiles. The histogram illustrates the distribution of both bead populations within the volume, plotted as a function of their lifetimes and provides a quantitative characterization of the fluorescence lifetime. The lifetime distributions had standard deviations of 0.22 ns (Nile Red) and 0.45 ns (yellow-green), at average photon counts of 10710 and 12330 per pixel, respectively. The histogram was generated from all pixels with fluorescence emission photon count greater than 8000. No Gaussian filtering was applied in this analysis. The volumetric and scan view of the fluorescent beads are presented in Supplementary Video 2 and Supplementary Video 3.

**Fig. 2.**
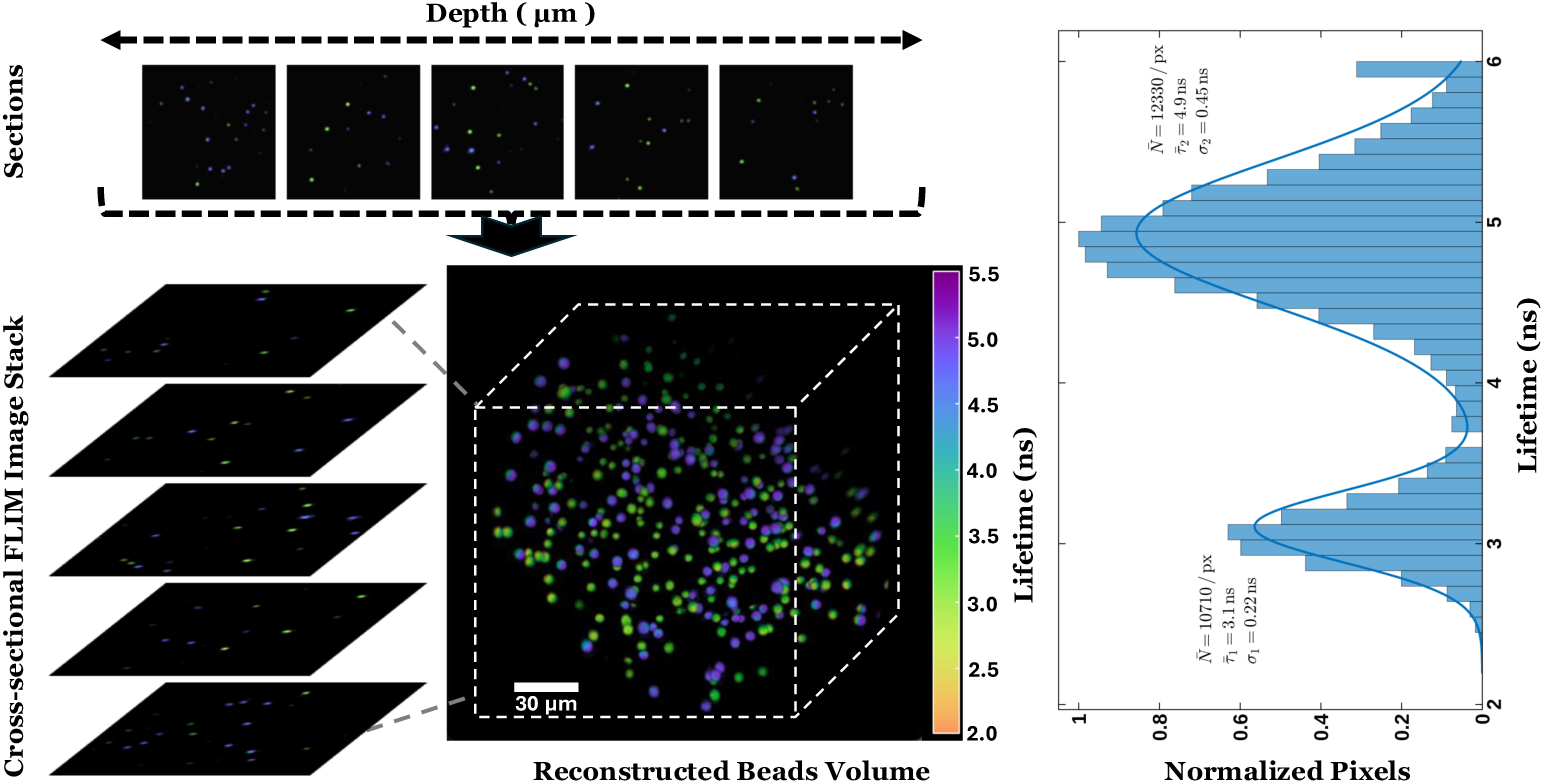
Volumetric fluorescence lifetime imaging of fluorescent beads: The mixture of yellow-green fluorescent beads (1.0 *µ*m) and Nile Red beads (1.0 *µ*m) was embedded in agarose gel. The mixture was confined within a rectangular capillary placed on a translational stage and illuminated using a two-photon scanning light sheet. Fluorescence emission was collected through a 10X/0.50 NA objective. As the sample stage was translated through the light sheet, sequential sectional images of the volume were acquired. These image slices were subsequently assembled into a three-dimensional reconstruction to visualize the full volumetric structure (middle panel). From this reconstructed dataset, we then computed a histogram (right panel) that represents the distribution of the two bead populations within the volume, plotted as a function of their lifetimes. The analyzed volume reveals average lifetimes (*τ̅*) of 4.9 ns for the yellow-green beads and 3.1 ns for the Nile Red beads. *N̅* and *σ* denote the average photons per pixel and corresponding standard deviation of lifetime. The complete volumetric and scan view is presented in Supplementary Video 2 and Supplementary Video 3.

### Quantitative comparison of fluorescence lifetime and intensity-based analysis

A fluorescent calibration slide (Thorlabs, FSK2) and a sample of mixed fluorescent beads (Nile Red and yellow-green, as above) were imaged to demonstrate that TPLS-EO-FLIM provides robust, intensity-independent contrast, suppresses intensity-based artifacts, and supplies complementary information that enables reliable discrimination of fluorophores that are indistinguishable by intensity alone. Samples were excited at 930 nm and emission was collected in the 530–570 nm band.

For the fluorescent calibration slide, line plots were extracted from the same spatial location in the intensity and lifetime images (Figure 3(a–b)) and analyzed to compare spatial behavior of intensity and lifetime across the field. The intensity line plot extracted from Figure 3(a-b) shows a gradual decrease in signal within the region *R*, while the corresponding lifetime trace in same region remains essentially uniform (Figure 3(c–d)). We observed that the intensity varied locally while the lifetime stayed nearly constant. This validates the capacity of TPLS-EO-FLIM to mitigate intensity-related distortions arising from uneven illumination, detector response, or local attenuation. To examine photon flux dependence, pixel-wise lifetime and intensity values were sampled along the identical lines *L*_1_ and *L*_2_ for multiple exposure times and plotted in Figure 3(e) (lifetime) and Figure 3(f) (intensity). Figure 3(g) summarizes these results by plotting mean fluorescence lifetime (left axis) and mean intensity (right axis) as functions of exposure time. While mean intensity changes with exposure, the mean lifetime remains largely stable.

**Fig. 3.**
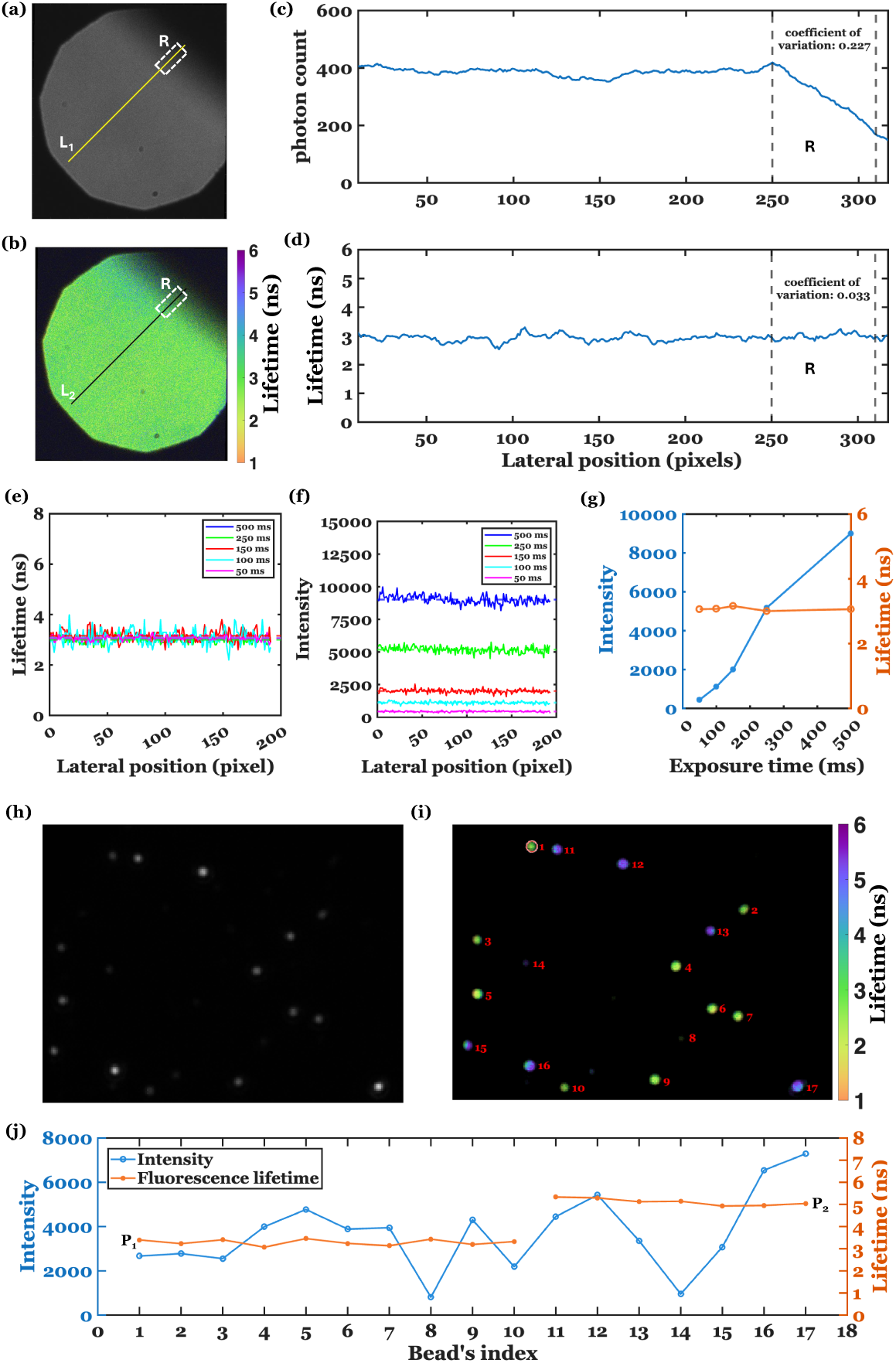
Quantitative comparison of fluorescence lifetime and fluorescence intensity measurements. A fluorescent slide and mixed fluorescent beads were excited at 930 nm, with emission collected within the 530–570 nm range. (a) The fluorescence intensity image of the slide. (b) The fluorescence lifetime image of the slide with overlaid intensity mask, intensity information mapped to opacity. (c) The intensity line plot across the line *L*_1_ in (a). (d) Fluorescence lifetime values across the line *L*_2_ in (b). (e) Fluorescence-lifetime profiles measured along the line *L*_2_ in (b) as a function of exposure time (colors correspond to exposure times shown in the legend). (f) The fluorescence intensity profiles along the line *L*_1_ in (a) for various exposure times. (g) Mean intensity and mean fluorescence lifetime across the lines *L*1 and *L*_2_ plotted as a function of exposure time. (h) Fluorescence intensity image of two populations of beads. (i) Fluorescence-lifetime image for the same field of view as (h); bead indices used in (j) are labeled on this image. (j) Mean intensity (left axis) and mean lifetime (right axis) for each bead, plotted against bead index. P1 and P2 represents the Nile red (*τ*_1_=3.1 ns) and Yellow-green (*τ*_2_=4.9 ns) bead populations in (i). The L_1_ and L_2_ lines are colocalized. The circle in panel (i), corresponding to index 1, represent the area for average intensity or lifetime calculation. The unit of intensity is the analog-to-digital unit (ADU) in (f), (g), and (j).

We extended the analysis to the bead samples. The intensity image (Figure 3(h)) does not distinguish the two types of bead population, whereas the lifetime map (Figure 3(i)) reveals the two distinct populations – Nile red (3.1 ns) and yellow-green (4.9 ns) beads. Bead indices were annotated on the lifetime map to preserve spatial correspondence for subsequent quantification. A per-bead scatter of average intensity versus average lifetime (Figure 3(j)) shows that beads with similar intensities can have distinct lifetimes and that beads with differing intensities can share similar lifetimes, demonstrating that lifetime contrast is largely independent of intensity and lifetime imaging is independent of intensity fluctuations caused by factors such as photon flux, excitation power or photo-bleaching.

### Dynamic fluorescence lifetime imaging of cardiac cycles of a zebrafish embryo’s heart

The capability of TPLS-EO-FLIM to image dynamic biological samples was demonstrated through imaging of cardiac cycles of a zebrafish embryo’s heart. The zebrafish embryo offers an established vertebrate model for cardiovascular research, enabling insights into mechanobiological processes such as cardiac function and hemodynamics during development. Figure 4 illustrates the cardiac dynamics of a 5 days post-fertilization (dpf) zebrafish embryo that carries a Tol2 transgene designed to express GFP in its heart under the control of the cardiac myosin light chain promoter (cmlc2:GFP). To ensure the tissue of embryo remains transparent during its developmental stages, the embryo was treated with N-Phenylthiourea (PTU), which effectively inhibits pigmentation. The zebrafish embryo was suspended in a thin-walled rectangular glass capillary tube filled with 1% agarose gel. This setup secures the embryo in place and minimizes spherical aberration during specimen scanning. Before suspension, the embryo was anesthetized with Syncaine to immobilize it and eliminate gross movement, allowing for stable and consistent imaging; cardiac contractions persist in these circumstances. This immobilization is essential for capturing the dynamic processes of cardiac activity, as it reduces artifacts caused by motion and ensures the integrity of fluorescence lifetime imaging data. On the other hand, the rapidity of the EO-FLIM technique offers the ability to capture lifetime dynamically, including motion on timescales slower than the frame rate, and further because both temporal gates are recorded simultaneously, motion between measurements does not result in motion artifacts as in FLIM techniques with single-photon counting or sequential gates. The specimen was excited at a wavelength of 930 nm, and the resulting fluorescence was collected through a 525 *±* 39 nm bandpass filter. A camera frame rate of 50 frames per second (20 frames per cardiac cycle) was used to record the cardiac cycles with minimal motional artifacts. The images were recorded using 4X (0.20 NA) objective lens. The recorded images were further analyzed, as discussed in the Methods section, to extract the spatial and temporal lifetime information and stacked in the time domain to make a movie (Supplementary Video 4) with spatial and dynamic fluorescence lifetime information. From these images, we estimated the biomechanical parameters, as detailed below. This methodology enables advanced imaging to study cardiac dynamics in zebrafish, providing insights into physiological mechanisms in a model organism widely used in developmental and cardiovascular research.

**Fig. 4.**
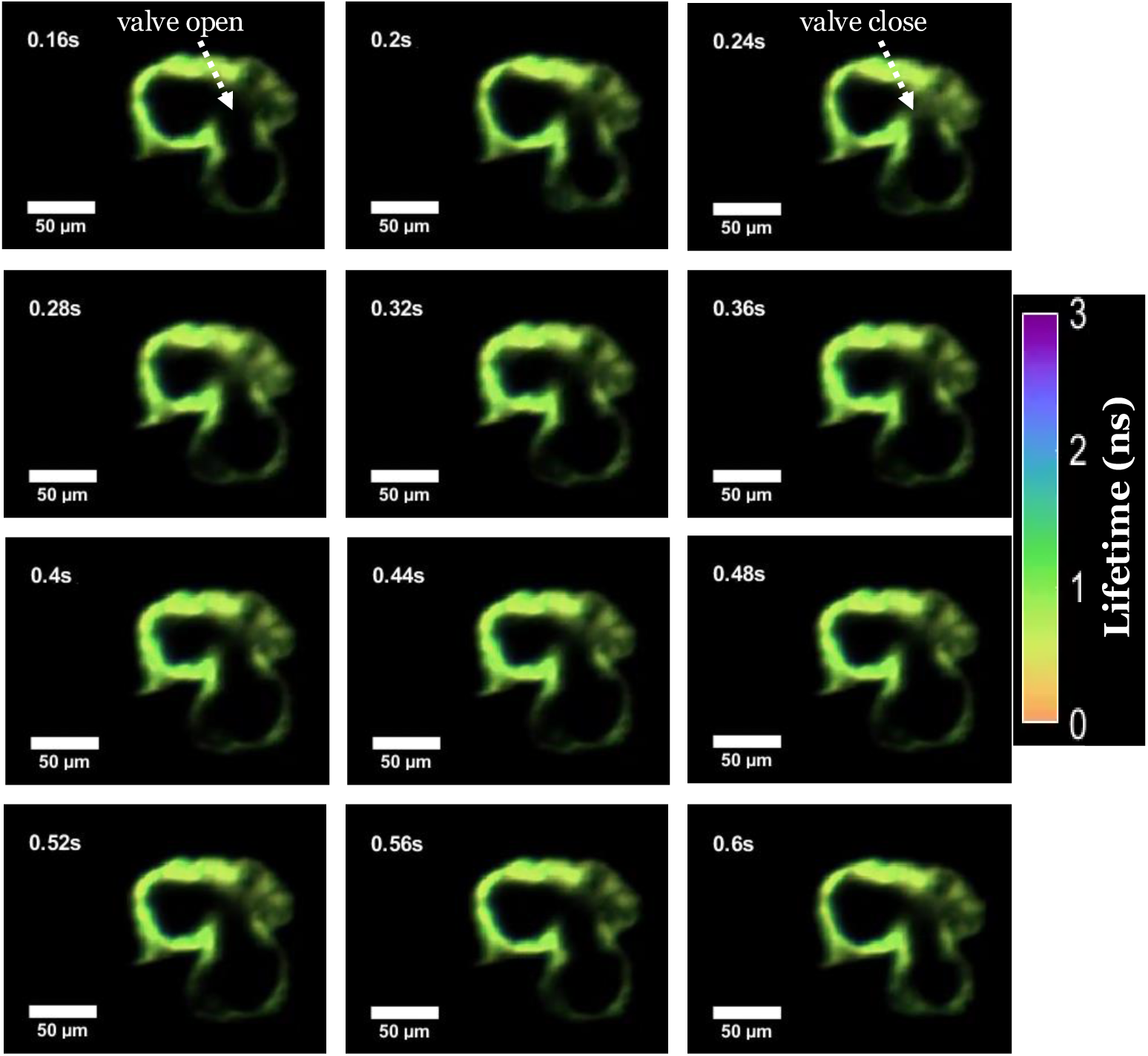
Cardiac dynamics of a zebrafish embryo: Time-lapse TPLS-EO-FLIM of a zebrafish heartbeat involves a 5-day postfertilization (dpf) zebrafish embryo with a genetically encoded GFP-expressing heart. Cardiac dynamics were captured at a high frame rate of 50 frames per second (every second frame shown). The recorded frames, at successive time points, illustrate the complete sequence of a single cardiac cycle, encompassing both the contraction (systole) and relaxation (diastole) phases of the heart, along with fluorescence lifetime information (Supplementary Video 4). The average frequency of a cardiac cycle was observed *∼* 2.5 Hz. Scale bar: 50 *µ*m

### Biomechanical parameter estimation of live zebrafish cardiac cycle

We show that TPLS-EO-FLIM can image at speeds fast enough to measure dynamic biomechanical parameters in living organisms. This establishes a foundation for future applications in which fluorescence lifetime measurements provide additional functional information beyond intensity alone. In particular, the system could be extended to investigate cellular stress, molecular interactions, and mechanical tension using FRET-based biosensors, where changes in donor fluorescence lifetime enable quantitative analysis through FRET-FLIM methodologies.

This capability was demonstrated through cross-sectional imaging of live zebrafish embryos. It was employed to characterize the dynamic behavior of the atrioventricular (AV) valve during diastole and systole (Figure 5(a) and Supplementary Video 4). For example, one of the primary metrics assessed was the temporal interval between valve states: the time difference between maximum (point 1) and minimum (point 2) valve opening (Δ*t*_12_) was measured at *∼* 80 ms, while the interval between minimum opening and the subsequent maximum (Δ*t*_21_) was *∼* 320 ms. The ratio (Δ*t*_21_)/(Δ*t*_12_) was found to be approximately 4.0 (*±*1, limited by the 20 ms frame interval), a value that remained stable over the next recorded cycles. This metric may be used to analyze physiological health of cardiac cycles. Figure 5(b) illustrates and quantifies the temporal variation in ventricular and atrial cross-sectional areas over the course of the cardiac cycle. Further time-lapse analysis of 2D cross-sectional images enabled quantification of additional biophysical parameters, including average pulse rate, maximum AV valve diameter, and Fractional Area Change (FAC). The average pulse rate was determined to be 2.5 Hz, calculated by measuring the time between two identical successive cardiac phases—either end-diastole (ED) or end-systole (ES)—as described by Hoage et al. [50]. The AV valve reached its maximum diameter of 25 *µ*m at a time stamp of 0.16 seconds. At 0.16 seconds, the atrial and ventricular areas (A*_a_* and A*_v_*) were measured to be 1836 *µ*m^2^ and 2828 *µ*m^2^, respectively. The value of FAC was 0.32 ± 0.01 (over three consecutive cardiac cycles), calculated from the end-diastolic (2926 *±* 78 *µ*m^2^) and end-systolic (1978 *±* 42 *µ*m^2^) ventricular areas. FAC is a well-established metric for evaluating ventricular contractility in zebrafish embryos, as previously reported by Haendchen et al. [51] and discussed by Yalcin et al. [52]. Additional cardiac parameters of interest — such as fractional shortening, ventricular volumes, stroke volume, ejection fraction and cardiac output — can be quantitatively estimated through the analysis of image stacks [51, 52].

**Fig. 5.**
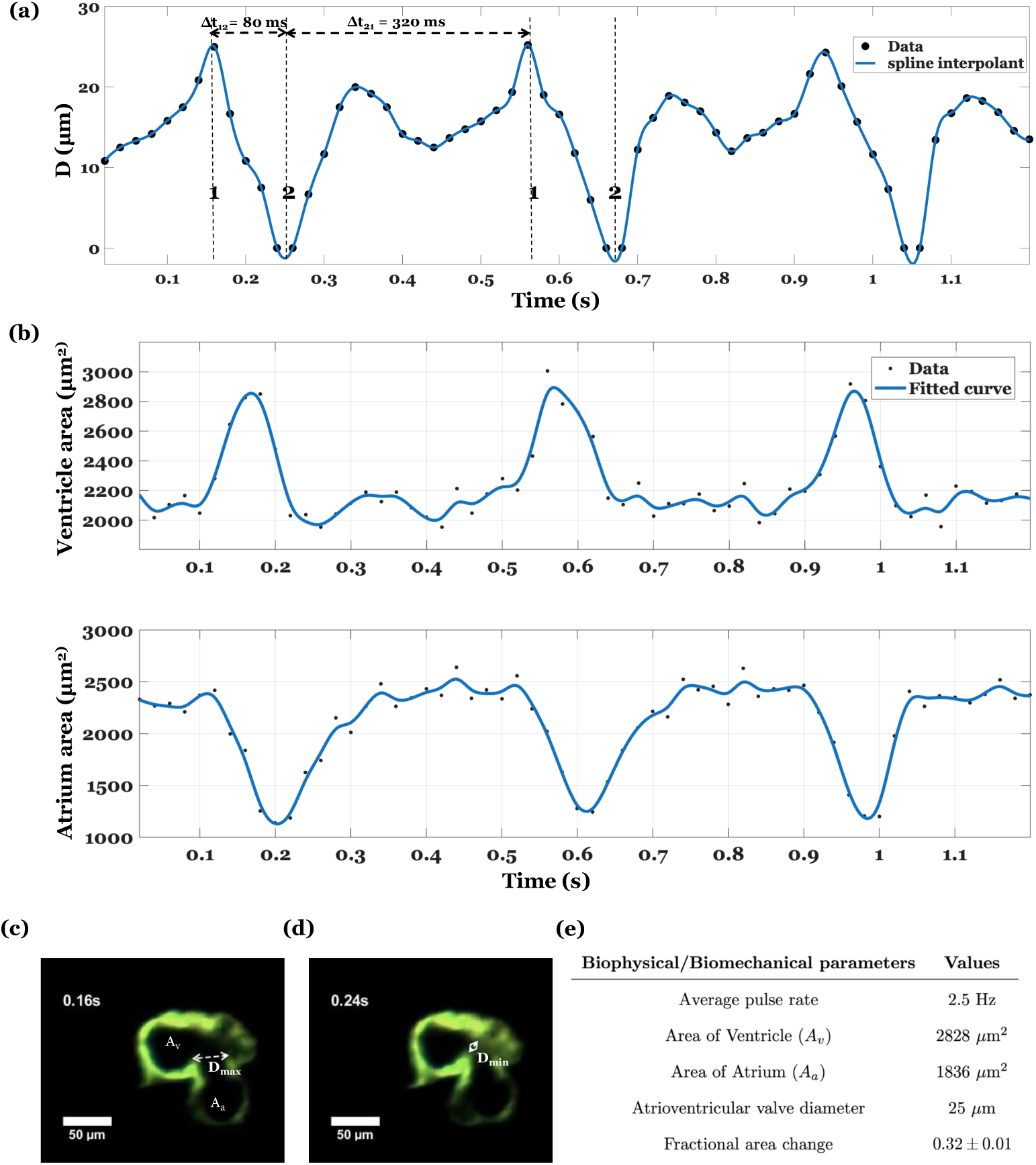
Biophysical parameter estimation associated with cardiac cycles: (a) The curve, atrio-ventricular (AV) valve diameter (D) evolution vs. time, shows the beating characteristics of the AV valve during diastole and systole periods. The time difference (Δ*t*_12_) between maximum valve opening (point 1) and minimum valve opening (point 2) is 80 ms, and the time difference between (Δ*t*_21_) Maximum opening and the next minimum opening is 320 ms. The ratio of (Δ*t*_21_) and (Δ*t*_12_) was found to be 4 and remained constant over the recorded cycles. (b) The temporal evolution of ventricular and atrium cross-sectional areas. (c and d) Cross-Sectional images at 0.16 s (diastole) and 0.24 s (systole). (e) The average pulse rate was found to be 2.5 Hz. The other biophysical parameters relevant to cardiac health were also calculated, such as area of atrium (A*a*), area of ventricle (A*v*) and the maximum diameter of the AV valve (D*max*). The values of A*a* and A*v* were calculated based on the frame captured at the 0.16-second time point (See Supplementary Video 4).

### Volumetric imaging of zebrafish embryo’s heart

We further illustrate cardiac imaging by demonstrating volumetric TPLS-EO-FLIM of the fixed zebrafish embryo’s heart. This work demonstrates the feasibility of using lifetime contrast for deep-tissue volumetric imaging of the entire embryonic heart, *∼* 125 microns into the tissue. The advantage of using lifetime contrast is that it avoids intensity artifacts as described above, and offers the possibility of using lifetime contrast to label structures (see below for a demonstration in plants). The same transgenic line and suspension protocols were used to record these images, as discussed in the previous section. The stage carrying the embryo was translated in steps of 2.5 *µ*m along the light sheet to record the sectional data of the GFP-expressing heart (cmlc2:GFP). The fluorescence was recorded with a 4x/0.20 NA objective. The exposure time of the camera was set to 250 ms, corresponding to 4 frames per second. A total of 100 frames were recorded to construct the complete volume, covering a depth of 250 *µ*m in *∼*25 s. Finally, the recorded frames were analyzed to extract the lifetime information, which was stacked together to visualize the volumetric TPLS-EO-FLIM of the embryo’s heart (Figure 6). The volumetric and cross-sectional scan views of the embryo’s heart are presented in Supplementary Video 5 and Supplementary Video 6.

**Fig. 6.**
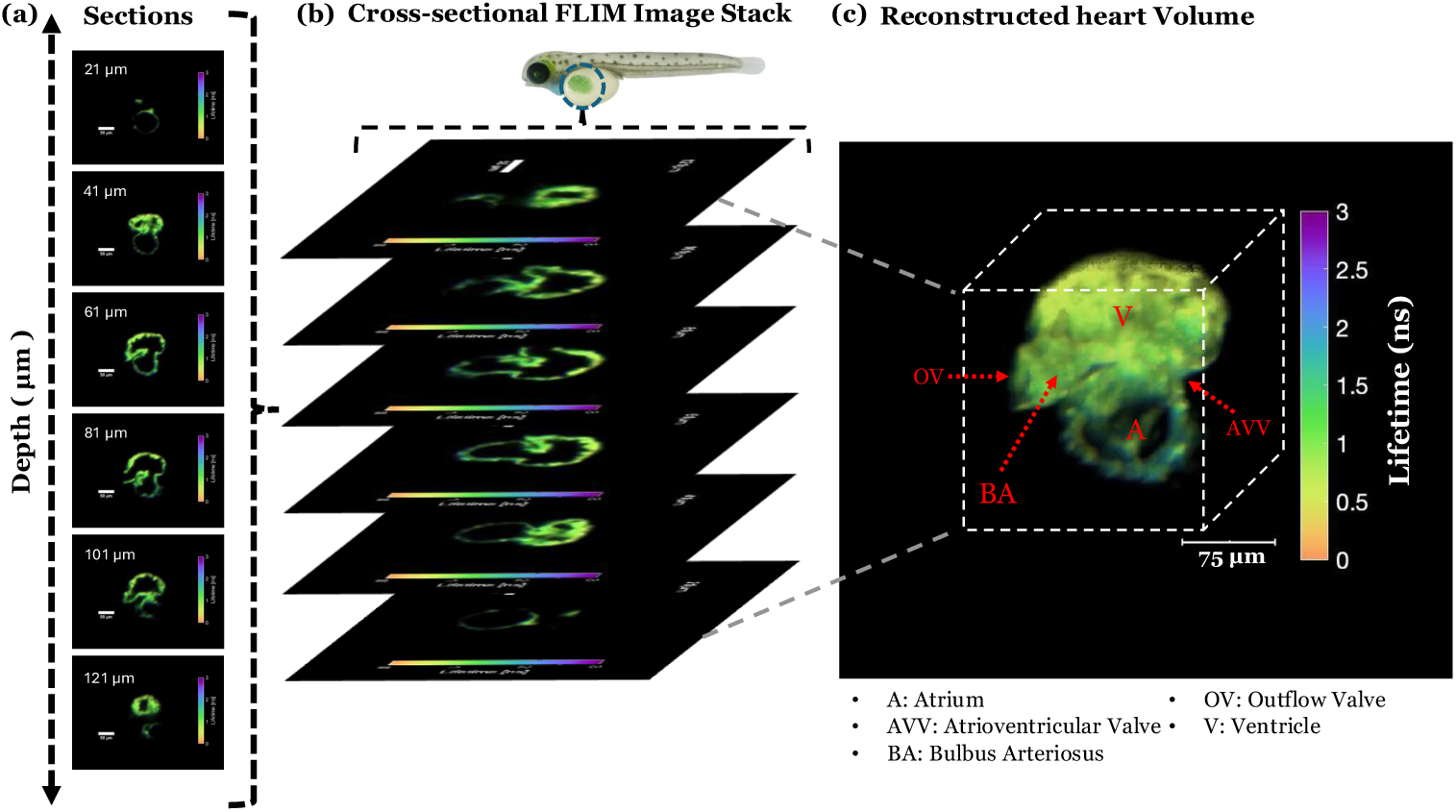
Volumetric TPLS-EO-FLIM of the zebrafish embryo’s heart: A GFP-expressing fixed embryo was translated in 2.5 *µ*m steps across the light sheet to acquire sectional data of its heart (a), with fluorescence collected through a 4x/0.20 NA objective. Imaging was performed at 250 ms exposure per frame. A total of 100 frames were recorded, analyzed for lifetime information, and stacked (b) to construct a complete volumetric FLIM visualization of the heart (c). Various regions of the organ are labeled with abbreviations and indicated by red arrows. A complete volumetric view and 2D cross-sectional scan of the embryonic heart are presented in Supplementary Video 5 and Supplementary Video 6, with additional anatomical details provided in Supplementary Information 4 through detailed cross-sectional view. Scale bar: 75 *µ*m

### Volumetric FLIM of *Arabidopsis thaliana* seedling root

TPLS-EO-FLIM can differentiate fluorescently tagged proteins and localize structures in a living sample by their lifetime. We demonstrated that TPLS-EO-FLIM can be used to obtain volumetric lifetime contrast by imaging *Arabidopsis thaliana* roots expressing multi-fluorescent proteins (multi-FP). Arabidopsis roots provide a stringent test of TPLS-EO-FLIM in living plant tissue, combining strong optical scattering with multiple co-expressed fluorescent proteins. We used this system to assess two aspects of performance: volumetric lifetime imaging of an intact root tissue, and lifetime-based discrimination of spectrally overlapping GFP- and YFP-tagged structures recorded at a single excitation wave-length and through a common emission filter rather than spectral separation. Here, we used the transgenic *Arabidopsis thaliana* line (Arabidopsis Biological Resource Center [ABRC], stock, CS16303), expressing CFP, GFP, YFP, and RFP markers designed to target nuclei, the plasma membrane, mitochondria, and plastids, respectively [53]. A 6-day-old specimen was immobilized in a sample holder. The holder was moved along the optical axis of the detection objective, allowing interaction with the light sheet. A 4x/0.20 NA detection objective collected the emitted fluorescence. A total of 125 frames were recorded at spatial intervals of 2.5 *µ*m, analyzed for fluorescence lifetime, and stacked to generate a volumetric reconstruction of the plant root (Figure 7). We specifically focused on the GFP- and YFP protein tagged structures. Control data were taken with selective GFP (520 *±*20) nm and selective YFP 590 *±* 20 nm filters, which independently confirmed structures, and are discussed in Supplementary Figure 6. We demonstrate that these structures can be distinguished based on their fluorescence lifetimes when excited at a common wavelength of 930 nm and detected through an emission filter with a band of 575 *±* 25 nm. In particular, we selected the filter range to capture signals from both the YFP and GFP spectra. The structures were then identified by the 2.3 *±* 0.25 ns average lifetime of the GFP and the 1.5 *±* 0.16 ns average decay lifetime of the YFP.

**Fig. 7.**
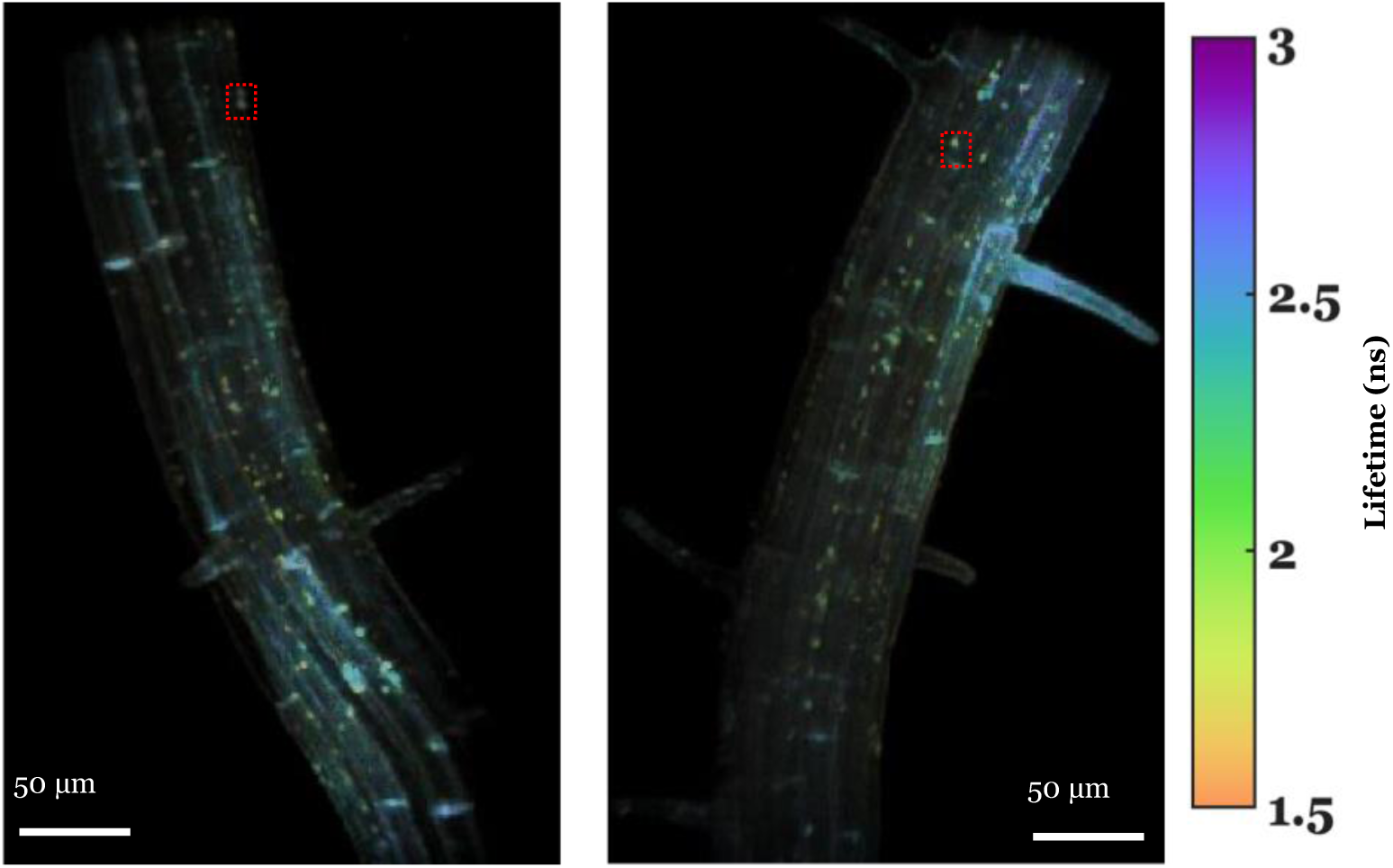
*Arabidopsis thaliana* root: Left and right panels show volumetric fluorescence lifetime images of selected regions of two different roots, recorded with TPLS-EO-FLIM. Multi-fluorescent protein (multi-FP) labeled seedlings were positioned on a custom-built sample holder and repositioned in the light sheet in 2.5*µ*m increments. A total of 125 frames were recorded, subsequently analyzed for spatial and fluorescence lifetime information and stacked to generate a volumetric visualization of the specimen (Supplementary Video 7 and Supplementary Video 8). Structures labeled with Cam53BD-GFP (plasma membrane marker) exhibit a longer fluorescence lifetime (2.3 *±* 0.25 ns) than those labeled with CoxIV-YFP (mitochondrial marker; 1.5 *±* 0.16 ns; red boxes)

## Discussion

TPLS-EO-FLIM introduces a high-speed approach to volumetric microscopy, facilitating the capture of deep structural and fluorescence lifetime information in diverse, living samples. Yellow-green and Nile Red fluorescent microspheres embedded in agarose gel were imaged with the imaging system. Distinct average lifetimes were resolved, confirming the system’s ability to differentiate fluorophores by their fluorescence decay profiles. We imaged a fluorescent slide and fluorescent beads to demonstrate that fluorescence lifetime remained stable despite variations in signal intensity across the field of view. This confirmed that TPLS-EO-FLIM effectively suppresses intensity-dependent distortions.

We demonstrate the utility of the method by imaging the live heart of a zebrafish embryo and live plant roots. We show that dynamic and full volumetric images can be obtained with lifetime contrast. This technique allowed for a detailed assessment of the structural and functional dynamics of these systems, demonstrating the utility of two-photon light sheet FLIM for whole-organ and high-resolution studies. The methodology aligns with recent technological advancements, incorporating selective plane illumination and electro-optic gating to enable rapid, phototoxicity-minimized fluorescence lifetime interrogation.

This system results in an average lifetime when two fluorophores colocalize. This limitation can be addressed by multispectral imaging, where signals are separated by two optical spectral filters, or phasor [28] or biexponential analysis. Here, the effective sectioning of the multiphoton excitation minimizes the presence of spatially distinct scatterers in an acquisition.

Future directions include its application in examining plant responses to environmental challenges, imaging complex organoid models in medical research, and integrating artificial intelligence to streamline the analysis of fluorescence lifetime data. Further refinement of this technique could further expand its applications in biology, agriculture, and medicine. Finally, the wide-field TPLS-EO-FLIM approach can be paired naturally with structured illumination to extract both super-resolution contrast and lifetime maps in a single acquisition [29].

## Methods

The system’s geometry follows the classic Selective Plane Illumination Microscopy (SPIM) configuration, where the illumination and detection subsystems are arranged orthogonally. It consists of three subsystems (Figure 1): (1) a digitally scanned two-photon light sheet illumination, (2) a sample platform, and (3) an EO-based wide-field detection subsystem. Each of these subsystems is described below.

### Digitally scanned two-photon light sheet illumination

For two-photon excitation, we utilized a femtosecond laser (Spectra-Physics Mai Tai HP Ti:sapphire ultrafast laser) with a repetition rate of 80±1 MHz, delivering 100 fs pulses and having a beam diameter (1*/e*^2^) of 1.2 mm. The repetition rate was reduced to 39 MHz using a pulse selection system to align with electro-optical wide-field detection requirements. This pulse selection system was comprised of an electro-optic modulator (Conoptics, Model 360-80), a synchronous countdown system (Conoptics, Model 305), and an amplifier (Conoptics, Model 25D).

The beam’s width was then expanded by a factor of two using a beam expander consisting of a pair of biconvex lenses and subsequently focused on the center of an axicon, followed by another biconvex lens to generate a non-diffractive Bessel beam. This Bessel beam was directed onto a galvano mirror (Thorlabs, GVSM002-EC) operating at 100 Hz to facilitate vertical scanning of the beam.

Following the scanning, the beam was passed through a scan lens (Thorlabs SL50-CLS2) and a tube lens (Thorlabs TTL200MP), which projected it on the back aperture of an objective lens (Thorlabs LMH-20X-1064, NA = 0.40, magnification = 20x, WD = 6.00 mm), thereby creating a tightly focused two-photon light sheet with a measured thickness of *∼*2.5 *µ*m. This scanned light sheet enabled precise two-photon excitation for thorough specimen interrogation.

### Sample Platform

The specimen holder was attached to the three-dimensional scanning stage, and the specimen was placed inside a square hollow glass tube. The rectangular geometry of the capillary tube plays a vital role in maintaining optical accuracy, ensuring that the scanned fluorescence data remain clear and well-resolved. Ensuring that the sample is precisely placed at the focal point of both the illumination and detection objectives (Figure 1), the sample volume can be scanned through linear translation stages along the X, Y, and Z axes. These stages are automated and facilitate control through MATLAB code and Kinesis Software.

### EO-based wide-field detection subsystem

The implementation of the detection subsystem, including resonant drive of the Pockels cell, is shown in Figure 1 and discussed in detail in [27]. The fluorescence light from the specimen was collected by the 4x objective (Thorlabs, USA, TL4X-SAP - 4X, NA= 0.20, WD = 17.00 mm) and 10x objective (Thorlabs, USA, TL10X-2P, NA= 0.50, WD = 7.77 mm) and split by a polarizing beam splitter into 50% s- and 50% p-polarization components. In this work, one polarized light component was passed through a single Pockels cell driven at 19.5 MHz, which rapidly rotates its polarization state as a function of applied voltage modulation through a resonant transformer. The operating mechanism of the Pockels cell is discussed by Bowman et al. [27]. The rotated light was then passed through a second polarizing beam splitter to create two gates (gated and ungated), and the light from these two gates was focused on the sCMOS camera (Hamamatsu, ORCA-Quest). The detection module can also be operated with two Pockels cells, one in each polarization path, to create a total of four gates; however, only the two-gate configuration was used in this study. The details of the implementation are discussed by Bowman et al. [25–27].

### Data Analysis

For data analysis, the two optical gates on the camera sensor were first registered with each other to have a pixel-to-pixel correspondence. The data were analyzed by creating a lookup table of an intensity gate ratio for each lifetime in the following manner. An instrument response function (IRF) was measured using quenched fluorescein, which emits near instantaneously (Supplementary Video 1 and Supplementary Information 3). The IRF effectively captures the convolution of the excitation and Pockels cell gating behavior. The lookup table ratios were computed by convolving the IRF with the lifetime exponential and then taking the ratio between the gated and ungated channels at the maximally sensitive IRF phase. That is, the fluorescence intensity trace while the Pockels cell cycles through phase delays can be understood as the convolution between the excitation function from the laser, gating function, and underlying exponential decay of the emitted fluorescence:

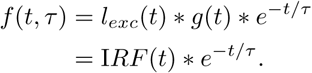

A ratio table or function can then be computed as

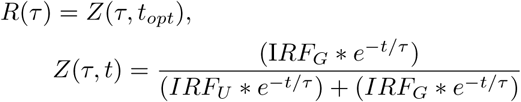

Here, I*RF_G_* and I*RF_U_* refer to gated IRF and ungated IRF, respectively. The delay time *t_opt_* is mapped to the Pockels cell phase defined relative to the laser excitation phase, and refers to the delay time at which the ratio between the gates changes most steeply as a function of lifetime during its oscillation, which is also when the ratio between the gates is maximized for an instantaneous emitter. The Pockels cell was set to this optimal phase to perform lifetime analysis using a single frame, as was done for most data including all videos, although data for stationary samples can also be taken and analyzed with full traces. At each pixel, the experimental ratio between the number of photons in each channel at the Pockels cell phase delay corresponding to *t_opt_* was matched to the simulated lifetime. Fluorescence lifetime can thus be determined using only one Pockels cell phase delay, and videos with short exposures can be taken where each frame can be analyzed for fluorescence lifetime.

After the lifetime image was generated, a transparency mask was applied to represent the intensity information. The lifetime maps were Gaussian-filtered using a nearest-neighbor approach that excludes pixels with lifetimes below 0.5 ns, suppressing unmatched-pixel artifacts during FOV registration and demonstrating the value of lifetime imaging in discriminating autofluorescence background signals. Three-dimensional rendering was performed in ImageJ and Napari [54].

### Sample Preparation

#### Fluorescent Beads

The 1% agarose gel matrix of the yellow-green (1.0 *µ*m, Excitation/Emission = 505/515 nm, Invitrogen, Thermofisher USA) and Nile Red (1.0 *µ*m, Excitation/Emission = 535/575 nm, Invitrogen, Thermofisher USA) fluorescent beads was prepared using agarose, beads, and water. One gram of ultra-pure agarose (Thermofisher Scientific) was dissolved in 100 mL of distilled water and heated to 121*^◦^*C. Once the solution became transparent, a syringe filter was used to filter it. Thereafter, the solution was cooled down to 50*^◦^*C, and then 5 *µ*L volumes of each fluorescent microsphere were mixed thoroughly in 2 mL of this solution. The mixture was left to cool to room temperature for 15 minutes to form a gel matrix.

#### Zebrafish lines and maintenance

All zebrafish experiments were conducted under protocols approved by the Stanford University Institutional Animal Care and Use Committee and conforming to appropriate city, state, and national regulations. Embryos and larvae were collected in E3 solution and treated with N-Phenylthiourea (PTU, 0.003%)(Sigma) from 23 hours post fertilization (hpf) to inhibit pigmentation. Prior to experimental procedures, embryos and larvae were anesthetized with 0.016% Syncaine (Syndel). The zebrafish line st167: Tg(cldnk:mCherry; cmlc2;GFP) carrying a Tol2 transgene using cardiac myosin light chain regulatory region to drive GFP expression (cmlc2:GFP) was used for fluorescence microscopy of heart tissue. Embryos and larvae were analyzed up to 5 dpf, before the onset of sexual differentiation.

#### *Arabidopsis thaliana* plants and maintenance

*Arabidopsis thaliana* multi-fluorescent protein (FP) (Arabidopsis Biological Resource Center [ABRC] stock CS16303) line was grown on 0.5 x Murashige-Skoog (MS) agarose plate (Caisson Lab-oratories, MSP06-50LT) with low-melting-point agarose (Thermo Fisher 16520100). This line expresses Gal4-CFP (nuclei), Cam53BD-GFP (plasma membrane), CoxIV-YFP (mitochondria), and RecA-RFP (plastids). The seeds were sterilized by 70% ethanol for 1 min, collected, washed in sterile water, suspended in 30% bleach for 30 min, and washed three more times in sterile water. Sterilized seeds were planted on the plate and maintained in a growth chamber (Percival AR-66L) at 50% humidity, 22*^◦^*C, and 100 *µ*mol/m^2^*/*s fluorescent illumination under a 10-h light/14-h dark cycle before imaging experiments. Seedling roots were imaged six days after planting.

#### Quenched fluorescein

A saturated solution of fluorescein was prepared in KOH water of pH 10. Thereafter, a 170 *µ*L volume of this solution was dissolved with 1 mL volume of saturated potassium iodide (KI) water to quench the fluorescence lifetime of fluorescein to the picosecond-scale lifetime. This mixed solution was used to record the instrument response function for the mid-vis spectral region (500-600 nm).

## Acknowledgments

We thank Dara Dowlatshahi for extensive discussion on samples and biological relevance of this imaging system. We thank the Department of Energy (DOE) office for funding support. This research was supported by the DOE Office of Science, Office of Biological and Environmental Research (BER), grant no. DE-SC0023076.

## Author Contributions

MK, AB, PK conceived the idea. PK performed the experiments. PK, JGK, DL prepared the samples. RK, PK analyzed and reconstructed the data. MK and SW supervised the research. MK, PK, RK, JGK, AB, DL, SW, MBM, and WT contributed to data interpretation and prepared the manuscript.

## Supplementary material

### S1. System configuration and implementation

**Fig. S1.**
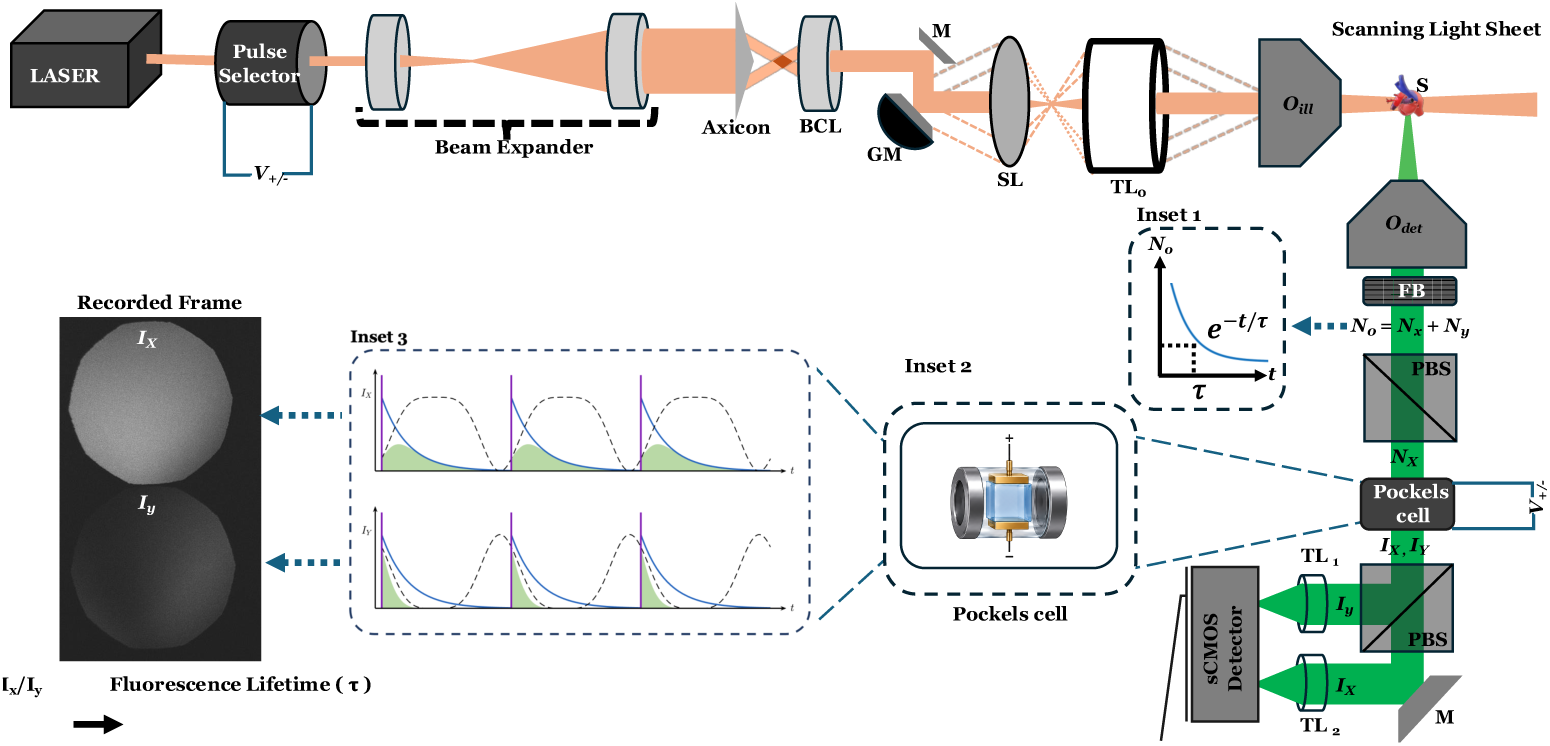
Electro-optic imaging based wide-field two-photon light sheet FLIM: The illumination and detection subsystems are arranged orthogonally for two-photon digitally scanning light sheet interrogation. A pulse laser operating at a 930 nm wavelength with a 39 MHz repetition rate produces a digitally scanned two-photon Bessel light sheet through an illumination objective (*O_ill_*) for specimen (S) sectioning. The specimen is translated in the XYZ directions for interrogation by the light sheet. Fluorescence emitted from the specimen is collected by a detection objective (*O_det_*), filtered through a filter box (FB) containing long-pass and notch filters, and then directed to a PBS-Pockels cell-PBS combination for polarization rotation as a function of time-varying applied voltage (V) on the Pockels cell (PC). The fluorescence light is split into its time-binned X and Y polarized components, which are then focused onto an sCMOS camera through a tube lens (TL) (see the recorded frame). The PC is synchronized with the laser repetition rate, which is 39 MHz. Inset 1 displays the fluorescence intensity decay over time, while Inset 3 presents a cartoon of the time-resolved split fluorescence intensity decay induced by modulation through the PC. Abbreviations: BCL, biconvex lens; GM, galvano mirror; *I_x_, I_y_*, intensity associated with x-polarized and y-polarized states; *N*_0_*, N_x_, N_y_*, photons associated with unpolarized, x-polarized, and y-polarized fluorescence; PBS, polarizing beam splitter; SL, scanning lens; TL, tube lens.

Fluorescence emission from the specimen is intrinsically isotropic, containing a distribution of all polarization states. Upon excitation, each fluorophore emits a photon burst whose total yield *N*_0_ can be expressed as the sum of its orthogonal polarization components, *N*_0_ = *N_x_* + *N_y_,* where *N_x_*and *N_y_*denote the x- and y-polarized photon counts, respectively. Critically, the intensity of this emission decays exponentially over time with a characteristic fluorescence lifetime *τ*, a parameter that is unique to each dye or protein and which forms the basis for lifetime-based contrast. Fig. S1 Inset 1 depicts the temporal decay curve of a representative fluorophore.

In the electro-optic fluorescence lifetime imaging microscopy (EO-FLIM) detection scheme, we manipulate light via polarization to temporally resolve fluorescence signals. To encode temporal decay into polarization, we first direct the emitted fluorescence through a polarizing beam splitter (PBS), separating it into s- and p-polarized beams. In this work, only one of the two beam paths passes through a single Pockels cell, an electro-optic crystal (lithium tantalate) whose birefringence is voltage-controlled; a second Pockels cell can be added in the other path to obtain four gates. A voltage driver applies a sinusoidal modulation at 19.5 MHz, synchronized precisely to the 39 MHz repetition rate of the Ti:sapphire laser. At each half-cycle of the applied voltage, the Pockels cell introduces a variable retardance that rotates the incoming s-polarization into two orthogonal states, hereafter labeled X and Y (Fig. S1 Inset 3). Thus, every point in the fluorescence decay is “time-stamped” by its induced polarization rotation.

Following modulation, a second PBS spatially separates the X- and Y-polarized components, which are then imaged through a tube lens onto an sCMOS detector array. These two images form the two gates (gated and ungated) used for lifetime analysis in the Methods section. The recorded intensity distributions, *I_x_*(*t*) and *I_y_*(*t*), correspond to photon populations leaving the Pockels cell at distinct phase delays. By computing the pixel-wise ratio *R* = *I_x_/I_y_,* and referencing a pre-calibrated mapping between R and *τ*, we reconstruct a high-resolution fluorescence lifetime map across the field of view as discussed in the methods section. This EO-FLIM approach leverages polarization modulation rather than time-gated detection, enabling continuous lifetime imaging at the full frame rate of the sCMOS sensor.

### S2. Excitation depth characterization using fluorescein

Setting up the optical sectioning parameters is essential to define the FOV, signal-to-background ratio, and axial resolution for an imaging setup. Also, proper characterization is required before the specimen’s volume interrogation. To measure the extent of the extent of the two-photon excitation, a stationary two-photon Bessel beam was propagated through fluorescein dye. The dye was contained in a 10 mm optical path length cuvette and placed at the intersection of illumination and detection objectives, so the resulting fluorescence was imaged from the side. A 20X illumination objective was used for illumination in this system, and a 4X detection objective was used to capture the excitation beam profile through fluorescence light emission (Figure S1). The FWHM of the line plot across the Bessel beam excitation region was found to be *∼* 340*µ*m.

**Fig. S2.**
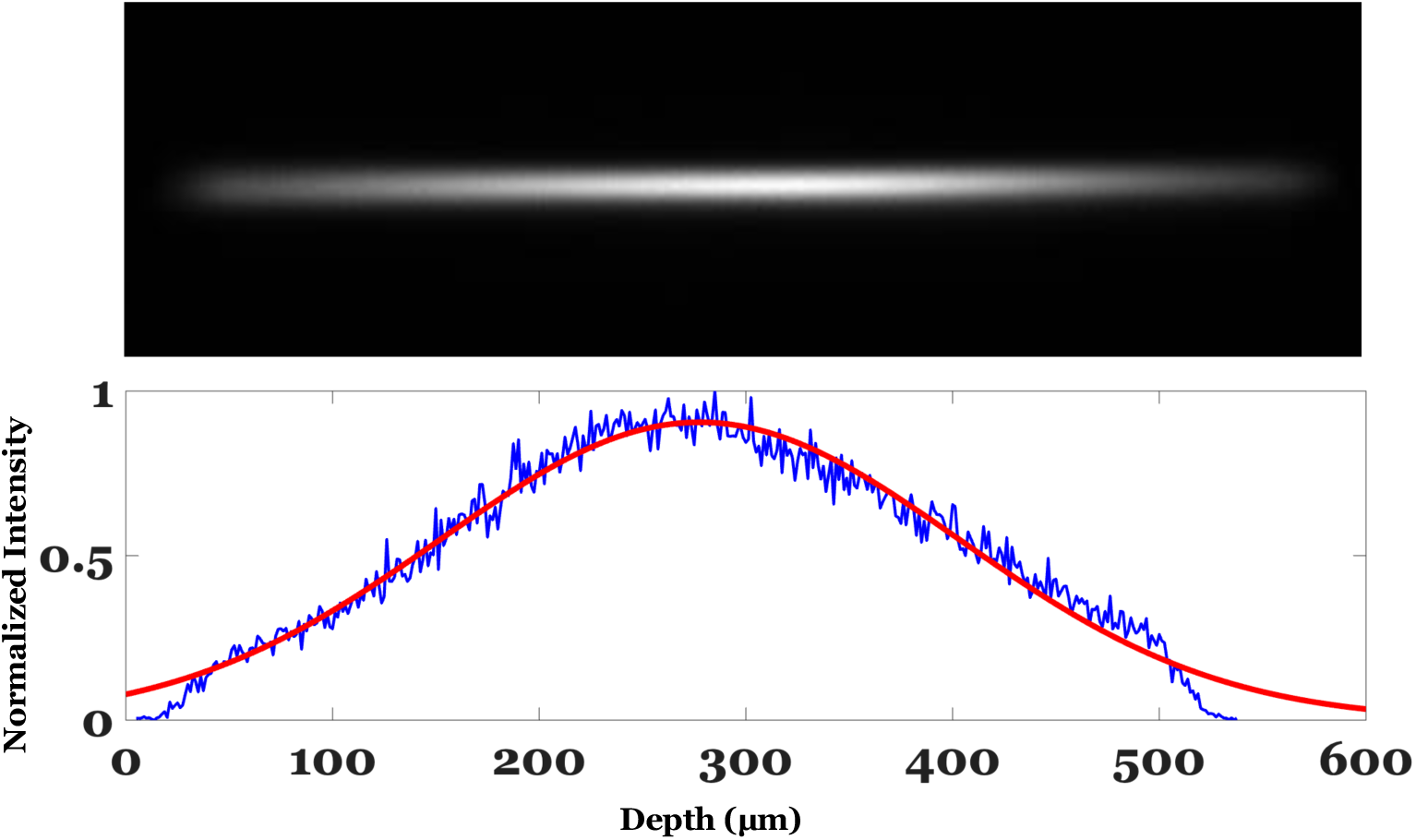
Light sheet depth characterization: The orthogonal view of the fluorescence emission from fluorescein along the optical path of a static two-photon Bessel beam quantifies the excitation depth of the imaging setup. The intensity plot across the excitation region shows a depth of *∼*340 *µ*m of the light sheet, as measured by the FWHM. The red line is a Gaussian fit to the measured intensity profile (in blue).

### S3. Instrument response function using quenched fluorescein

The instrument response function (IRF) of our EO-FLIM setup was calibrated using a potassium-iodide-quenched fluorescein solution. Native fluorescein exhibits a fluorescence lifetime on the order of nanoseconds, but the addition of KI — a heavy-atom quencher that enhances inter-system crossing — shortens its decay to approximately picoseconds scale. This ultrafast, near-impulsive fluorescence transient serves as an effective reference for deconvolving and characterizing the IRF of the detection system.

**Fig. S3.**
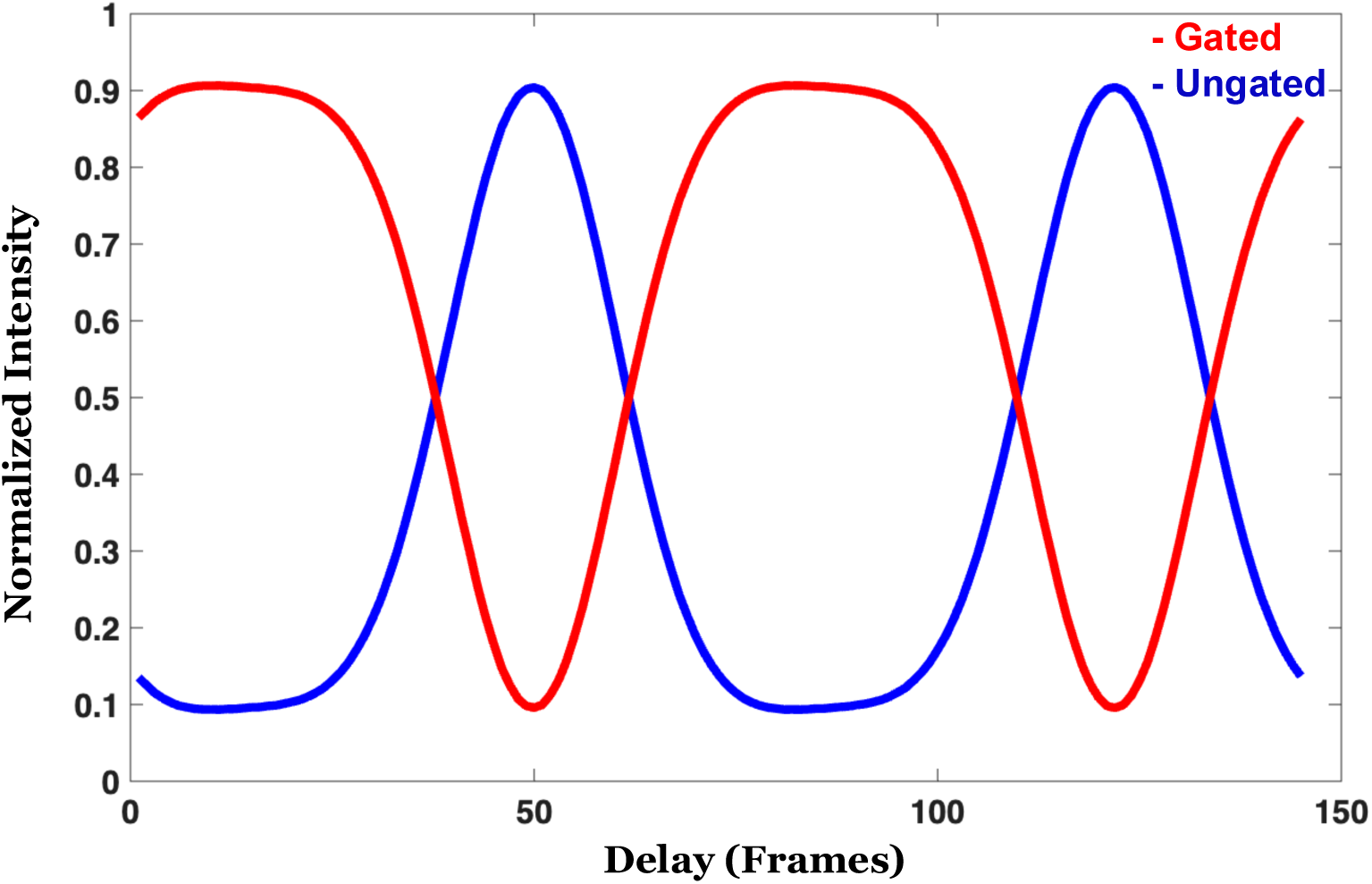
Instrument response function using quenched fluorescein: Normalized intensities from gated and ungated images are plotted as a function of phase delay to analyze temporal modulation.

### S4. Anatomical details of the zebrafish embryo’s heart

**Fig. S4.**
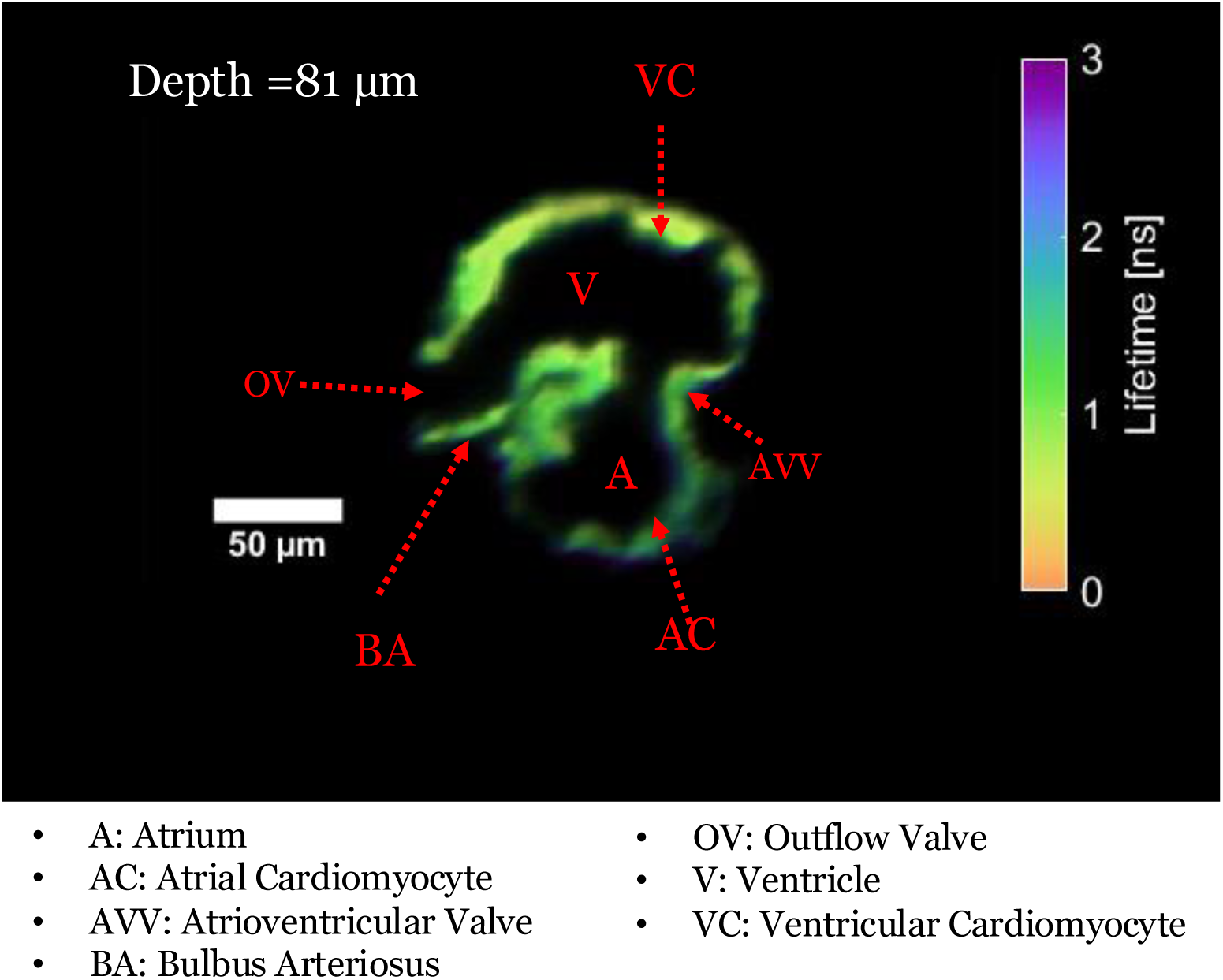
Cross-sectional FLIM image of 5 dpf zebrafish embryo’s heart: Anatomical details of the zebrafish embryo’s heart:

### S5. Lateral resolution characterization of the system using fluorescent beads

To assess the lateral resolution of the imaging system, microspheres with a diameter of 1.0 *µ*m were employed as test samples. These beads act as near point sources when captured by the instrument, making them suitable for evaluating the system’s point spread function (PSF). By analyzing the recorded bead images, the PSF was measured and found to have a width of approximately 1.96 *µ*m. This measurement reflects the effective lateral resolution of the system, indicating the smallest structural detail that can be reliably distinguished in the acquired images.

**Fig. S5.**
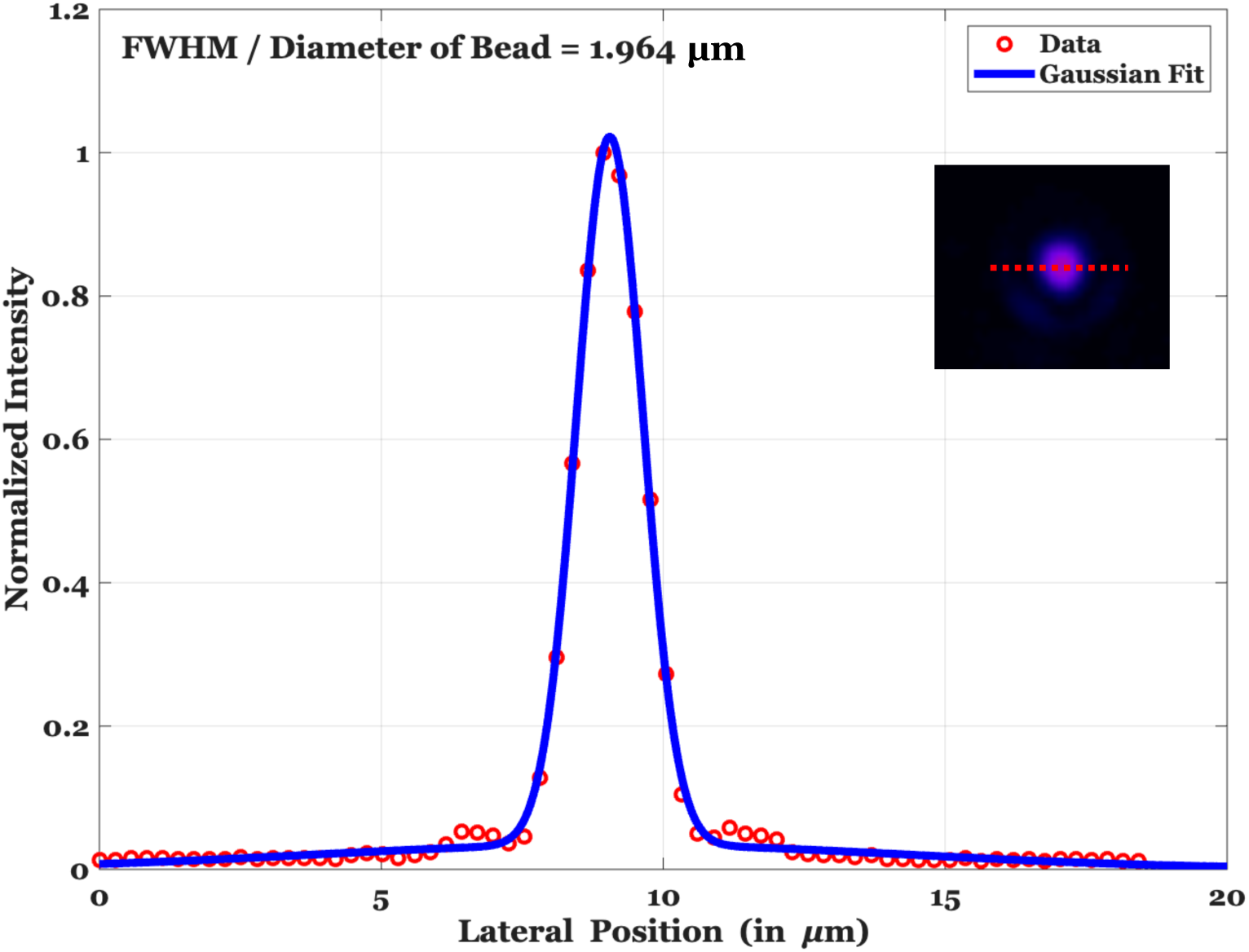
Characterization of the system’s lateral resolution using microspheres: The beads act as point-like emitters when imaged, allowing the point spread function (PSF) to be measured. Analysis of the recorded 1.0 *µ*m bead profiles yielded a PSF of full width at half maximum (FWHM) of approximately 1.96 *µ*m, which defines the effective lateral resolution of the imaging system.

### S6. TPLS-EO-FLIM of *Arabidopsis thaliana* root

**Fig. S6.**
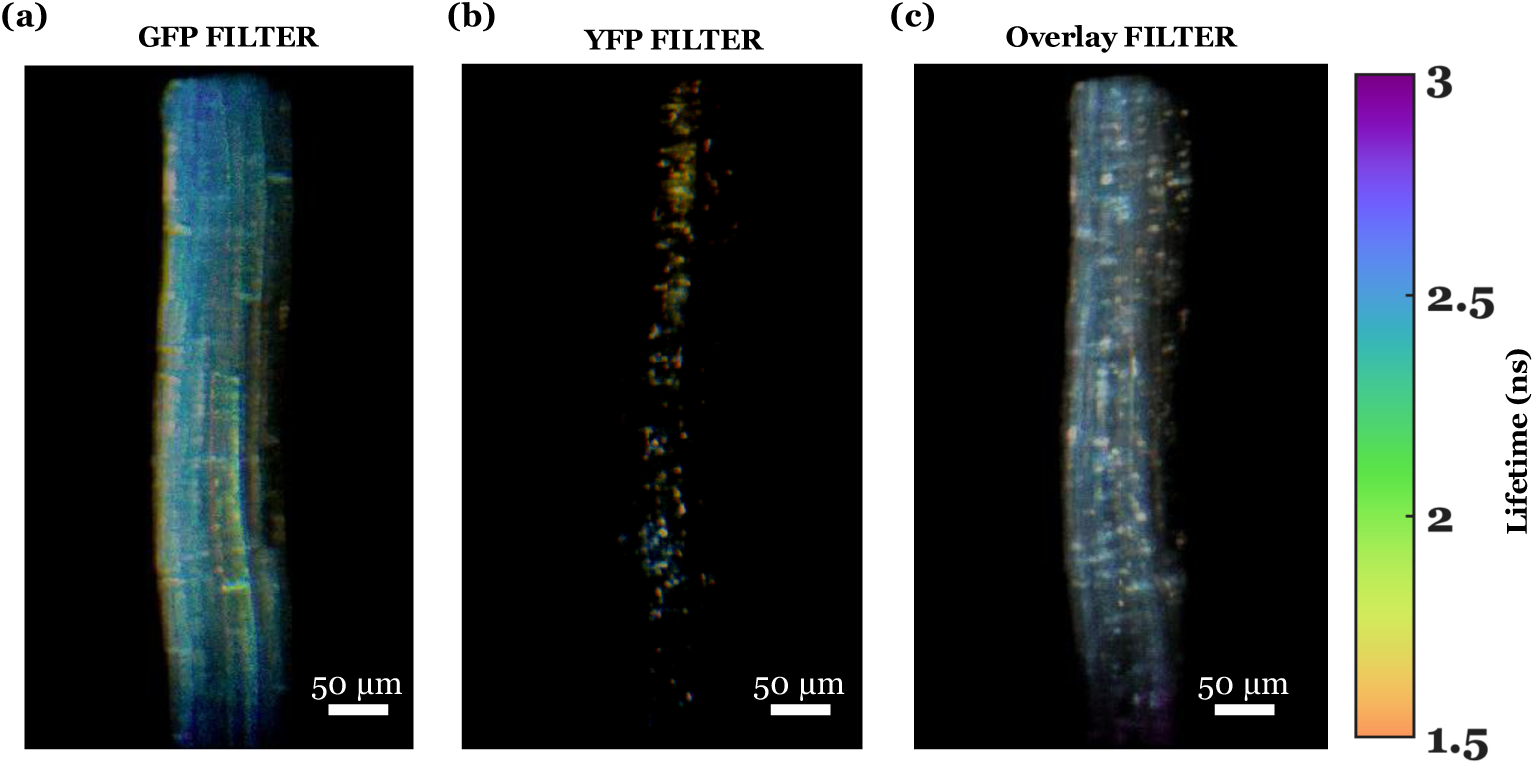
GFP and YFP fluorescence lifetime imaging of *Arabidopsis thaliana* root: Six-day-old multi-FP seedlings were excited at 930 nm. (a) GFP (Cam53BD-GFP) fluorescence lifetime image acquired from the *Arabidopsis thaliana* root through the GFP filter (520 *±* 20 nm). (b) YFP (CoxIV-YFP) fluorescence lifetime image acquired from the same root region through the YFP filter (590 *±* 20 nm). (c) Composite fluorescence lifetime image acquired through the wide range band pass filter (550-600 nm), showing the combined GFP and YFP lifetime signals and illustrating their relative spatial distribution and lifetime contrast within the root tissue. The colorbar represents the fluorescence lifetime in ns.

## Supplementary Videos details

Video 1: Instrument response function and Pockels cell modulation.

Video 2: Volumetric scan of the fluorescent beads.

Video 3: Volumetric view of the fluorescent beads.

Video 4: Cardiac dynamics of zebrafish embryo’s heartbeat.

Video 5: Volumetric view of the zebrafish embryo’s heart.

Video 6: Volumetric scan of the zebrafish embryo’s heart.

Video 7: Volumetric view of the *Arabidopsis thaliana* root 1.

Video 8: Volumetric view of the *Arabidopsis thaliana* root 2.

